# Circulating miRNA–Protein Signatures Predict Outcomes in Pediatric Dilated Cardiomyopathy

**DOI:** 10.64898/2026.03.17.712519

**Authors:** Amanda R. Vicentino, Anis Karimpour-Fard, Taye Hamza, Brian L. Stauffer, Kory J. Lavine, Shelley D. Miyamoto, Steven Lipshultz, Carmen C. Sucharov, Pediatric Cardiomyopathy Registry Investigators

**Affiliations:** Department of Medicine, Division of Cardiology, University of Colorado Anschutz Medical Campus CO; Department of Medicine, Division of General Internal Medicine, University of Colorado School of Medicine CO; Carelon Research, Newton Office, MA; Department of Medicine, Washington University in St. Louis MO; Department of Pediatrics, Division of Cardiology, University of Colorado Anschutz Medical Campus CO; Department of Pediatrics, University at Buffalo Jacobs School of Medicine and Biomedical Sciences NY

**Keywords:** pediatric dilated cardiomyopathy, secreted factors, biomarkers, machine learning

## Abstract

**Background:** Pediatric dilated cardiomyopathy (DCM) is a rare, progressive heart disease with variable outcomes that range from recovery to heart transplantation. To date, there are no prognostic biomarkers for children with DCM. Identifying circulating biomarkers that are associated with clinical outcomes is critical for personalized management.

**Methods:** miRNAs were identified by RNA-seq, whereas proteins were identified by SomaScan^®^. Machine learning methodologies were used to explore the predictive ability of circulating factors identified from serum samples collected at the time of presentation with acute heart failure.

**Results:** Thirty patients experienced poor outcomes (cardiac transplantation, mechanical circulatory support, or death) and 19 patients recovered left ventricular function. Distinct miRNA and protein signatures differentiated outcomes groups. Top candidate proteins (COL2A1, CXCL12, and ADGRF5) and miRNAs (miR-874-3p, miR-335-3p, miR-323a-3p) demonstrated strong discriminatory performance within the study cohort (recovered vs poor outcomes; Area Under the Curve of 0.92). Ingenuity Pathway Analysis implicates cardiac remodeling, fibrosis, and inflammatory signaling as central pathways differentiating patient outcomes.

**Conclusions:** Circulating miRNA and protein signatures at presentation identify a circulating molecular signature associated with divergent clinical trajectories in pediatric DCM. These findings support the potential utility of multi-omic biomarkers for early risk stratification and provide insight into mechanisms underlying divergent outcomes.

**CLINICAL PERSPECTIVE:** **What Is New?**

- Circulating miRNA and protein profiles measured at presentation distinguish children with pediatric DCM who recover from those who progress to advanced heart failure.
- A combined multi-omic biomarker demonstrated strong discriminatory performance in this cohort (AUC 0.92).
- Pathway analysis implicates extracellular matrix remodeling, fibrosis, and inflammatory signaling in children with adverse clinical trajectories.

**What Are the Clinical Implications?**

- Serum-based molecular biomarkers may enable earlier risk stratification in children presenting with dilated cardiomyopathy.
- Multi-omic integration may improve identification of pediatric patients at risk for transplantation, mechanical circulatory support, or death.
- These findings support further validation of circulating biomarker panels to guide personalized management in this rare disease.

**RESEARCH PERSPECTIVE:** **What New Question Does This Study Raise?**

- Can integrated circulating miRNA–protein signatures identify biologically distinct trajectories of recovery versus progression in children with dilated cardiomyopathy?
- Do circulating molecular profiles reflect underlying disease mechanisms that determine divergent clinical outcomes in pediatric DCM?

**What Question Should Be Addressed Next?**

- Do the pathways identified by integrated miRNA–protein analysis (fibrosis, remodeling, and inflammation) play causal roles in determining recovery versus progression?
- Can multi-omic biomarkers be incorporated into prospective studies to improve early risk stratification and guide clinical management?

## INTRODUCTION

Dilated cardiomyopathy (DCM) is the most common cause of heart failure and the leading indication for cardiac transplantation in the pediatric population over the age of one. Although rare, pediatric cardiomyopathies are associated with poor outcomes as nearly 40% of children who present with symptomatic cardiomyopathy either undergo heart transplantation or die within the first 2-5 years after diagnosis.^1–5^ The causes of pediatric DCM are heterogeneous, ranging from genetic variations that affect myocardial processes to systemic diseases that result in diffuse myocardial injury.^5^ However, in approximately two-thirds of cases, the cause remains undetermined, making idiopathic DCM the most common diagnosis.^6^

Although the majority of children with DCM has poor outcomes, recovery of cardiac function has been reported in 15-35% of cases across multiple single- and multi-center studies.^4,7,8^ While a uniform definition of recovery is still needed, myocardial recovery is generally defined as normalization of left ventricular (LV) size and function. Younger age, less LV dilation, and higher LV fractional shortening or ejection fraction have been the most consistent predictors of recovery in studies where this information was available.^8,9^

A few studies have identified potential biomarkers among children who eventually recover ventricular function. Fenton et al. reported age, LV end-diastolic diameter (LVEDD) z-score and N-terminal prohormone of BNP (NT-pro-BNP) levels were associated with outcomes. However, LVEDD z-score was not a predictor of outcomes, whereas older age and higher NT-pro-BNP levels were associated with poor outcomes.^8,9^ In contrast, Everitt, et al. showed that a lower LVEDD z-score and younger age were associated with a higher probability of recovery. Yet, to date, a strong biomarker of outcomes in children with DCM has not been established.^8^

Our group has explored the secretome as a potential biological milieu of factors that may support the diagnosis and prognosis of heart failure. The secretome encompasses proteins, microRNAs, and vesicles secreted by the heart or other organs.^10,11^

Secreted microRNAs (miRNAs, miRs) are small noncoding RNAs of ∼22 nucleotides detectable in blood that have emerged as promising diagnostic biomarkers. Our group has consistently demonstrated that miRNAs play a crucial role in improving our understanding of biomarkers and mechanisms underlying cardiovascular diseases, including hypertrophic cardiomyopathy and pediatric DCM.^12–15^ We previously showed, through miRNA arrays, that the circulating miRNA profile of children with severe heart failure secondary to DCM serves as a biomarker of outcome, distinguishing children with the potential for recovery from those who require transplantation.^12^ We also identified miRNA signatures in the myocardium of children with DCM, that were further stratified by age and sex.^13^ Furthermore, we investigated a relationship between serum and cardiac miRNAs.^14^

Secreted proteins such as brain-type natriuretic peptide (BNP), NT-proBNP, and cardiac troponins have been incorporated into the guidelines for adult heart failure diagnosis and management by both the European Society of Cardiology (ESC)^16^ and the American Heart Association (AHA).^17^ Additional diagnostic biomarkers, including markers of inflammation (e.g., soluble ST2 receptor), oxidative stress (e.g., growth differentiation factor-15), and cardiac remodeling (e.g., galectin-3), have also been proposed to guide heart failure therapy.^17^ Importantly, a comparison of circulating proteins in adult and pediatric DCM identified distinct biomarker profiles between the two populations, supporting the concept that adult and pediatric DCM are distinct diseases.^18^ Nevertheless, because pediatric DCM differs significantly from adult DCM, no specific protein biomarkers have been validated in children.

The potential for recovery underscores the importance of identifying reliable predictive biomarkers in children with DCM. Accurate predictors at the time of acute heart failure could help avoid unnecessary cardiac transplants in patients with the potential for recovery. In this work, we expanded on our prior investigations and explored circulating miRNAs, identified by RNA-seq, and circulating proteins, identified by SomaScan^®^, as prognostic biomarkers of recovery in children with DCM. Furthermore, we examined whether integrating miRNA and protein signatures enhanced discriminatory performance for DCM outcomes.

## METHODS

### Blood Collection

Human subjects were boys and girls of all races and ethnic backgrounds, < 21 years old, with a diagnosis of DCM. Blood was collected at the time of presentation with acute heart failure at Children’s Hospital Colorado and the Pediatric Cardiomyopathy Registry (PCMR)-associated centers, and at least six months after the first blood draw or when the patient met an outcome. Inclusion criteria were age < 21 years and echocardiographic or cardiac MRI evidence of DCM, defined as having at least 2 qualifying measurements present at the time of diagnosis including an LV ejection fraction (LV EF) or an LV fractional shortening (LV FS) > 2 s.d. below the normal mean for age, LV end-diastolic thickness-to-dimension ratio < 0.12, and LV end-diastolic dimension (LV EDD) or volume > 2 s.d. above the normal mean for BSA. Exclusion criteria included a prior transplant, other functional types of cardiomyopathy (e.g., restrictive, LVnon-compaction) and secondary cardiomyopathies (e.g. anthracycline toxicity, congenital heart disease).^19^

The PCMR study is a prospective cohort study of pediatric cases of DCM conducted at 11 pediatric cardiology centers in the US and Canada utilizing the established network of clinical center and infrastructure of the NHLBI-funded Pediatric Cardiomyopathy Registry. 150 patients with DCM were targeted for enrollment over 18 months with up to 24 months of follow-up. Subjects were < 21 years old. Eligibility included echocardiographic criteria according to the data collection schedule. The date of cardiomyopathy diagnosis is the date of the earliest available echo or MRI image or report that confirms the diagnosis of cardiomyopathy.

Following completion of clinical studies, leftover banked serum was stored at –80°C. Sera used for this study was approved by the Institutional Review Board from the collection hospitals/institutions. Detailed clinical characteristics for all patients are listed in Table S1.

### miRNA Extraction and Sequencing

miRNA was sequenced by LC Sciences (Houston, TX). All extracted RNA was used for library preparation following Illumina TruSeq Small RNA protocols (Illumina, San Diego, CA, USA). Library quality and quantification were assessed using an Agilent 2100 Bioanalyzer High Sensitivity DNA Chip. Single-end 50 bp sequencing was performed on the Illumina HiSeq 2500 system according to the manufacturer’s protocols. Copy number normalization used a modified global normalization based on a subset of sequences consistent across samples. As sera volume from pediatric patients is relatively small, miRNA was extracted from 35 µL of human sera. To establish if this sufficiently capture miRNAs present in serum, three sera volumes (200 µL, 100 µL, and 35 µL) from two patients were evaluated by RNA-sequencing. These yielded a within-sample R value of 0.99 (data not shown).

### miRNA Sequencing

RNA sequencing reads were obtained using the Illumina HiSeq 2500. Read quality was checked using FastQC (http://www.bioinformatics.babraham.ac.uk/projects/fastqc/). Adapter sequences were trimmed from raw sequencing files, and the reads were aligned to miRBase 22.0 using exact-match alignment to determine genomic positions. The number of reads mapping to each miRNA was quantified, and counts were normalized using edgeR^20^

### SOMAscan Analysis

Sera (55 µL) were used for SOMAscan^®^ analysis, which detected 5,116 aptamers. Protein integrity was evaluated using established QC metrics. Briefly, samples were mixed with beads conjugated to 5,116 validated aptamer probes. Protein-aptamer complexes were purified, and bound aptamers were quantified on a custom Agilent microarray. Raw signal intensity data were uploaded to the SOMAscan^®^ server for QC and standardization. The final dataset included standardized relative fluorescent units (RFU) for 5,116 proteins, SOMAscan^®^ Quality Statements (SQS), and basic differential expression. The assay output included both the identity and quantity of aptamer-bound proteins reflecting serum protein concentration.

### Proteomics

Somascan data were imported from a tab-delimited file using R. The dataset was filtered to remove unwanted sample types, and sample IDs were processed to ensure uniqueness. The annotation data, including SomaID, UniProt and biomarker targets, were matched with the corresponding sample data. The final dataset, combining both the biomarker data and annotations, was exported as a CSV file for downstream analysis.

### Integration of miRNA and proteomics

To integrate protein and miRNA expression data, we applied a rank-based normalization strategy to each dataset independently. For each sample (column), feature values were converted to ranks, and the ranks were divided by the total number of features to obtain normalized values between 0 and 1. This approach minimizes platform-specific scale differences while preserving the relative ordering of features within each sample. After normalization, the ranked protein and miRNA matrices were combined into a single integrated dataset, generating one file containing both data types for downstream analyses.

### miRNA and Protein Classification Analysis

Classification analyses were performed separately for miRNAs and proteins, as well as using a combined dataset integrating both biomarker types. The Random Forest (RF) algorithm implemented in the Random Forest package in R version 4.3.3 was used to build classification models, with 50,000 trees grown for each model. The primary objective was to identify circulating miRNAs and/or proteins that differentiate clinical groups.

Two complementary analytical approaches were applied: a machine-learning–based method (Random Forest) and traditional statistical methods (logistic regression and Wilcoxon rank-sum test). For the RF models, feature importance measures were used to rank candidate biomarkers within each dataset (miRNA-only, protein-only, and combined).

In addition, hierarchical clustering analysis was conducted using the top-ranked features from each analysis (miRNA-only, protein-only, and combined) to validate and visualize similarities and grouping patterns among samples and to assess the discriminative performance of the identified biomarkers.

### Model Evaluation and Statistical Analysis

All downstream analyses were performed separately for the miRNA-only dataset, the protein-only dataset, and the combined miRNA–protein dataset.

Random Forest (RF) results were visualized using multidimensional scaling (MDS) plots derived from proximity matrices (dimension 1, x-axis; dimension 2, y-axis). Variable importance was assessed using mean decrease in accuracy and mean decrease in Gini index, and corresponding importance plots are presented in the Figures. We selected the top three miRNAs and proteins for subsequent analyzes as they represent the most influential variables explaining the outcome while maintaining model interpretability and robustness. In our analysis, these top-ranked factors accounted for a disproportionate share of total variable importance, with subsequent variables contributing marginal gains. Including additional lower-ranked predictors did not meaningfully improve model performance, suggesting diminishing returns beyond the top three features.

Hierarchical clustering was conducted using the *hclust* function in R (Ward’s minimum variance method) and the ClassDiscovery package to evaluate sample grouping based on top-ranked features.

Wilcoxon rank-sum tests were applied to identify differentiating miRNAs and proteins. Multiple comparisons were adjusted using false discovery rate correction, and features with q-values ≤ 0.1 were considered statistically significant. Fold change (log_2_ scale) was calculated as the difference between group means. Wilcoxon rank-sum tests were applied to identify differentiating miRNAs and proteins. Multiple comparisons were adjusted using false discovery rate correction, and features with q-values ≤ 0.1 were considered statistically significant. Fold change (log_2_ scale) was calculated as the difference between group means. In addition, logistic regression analyses were performed to evaluate the association between individual features and the outcome. Odds ratios (ORs) with corresponding p-values were estimated to quantify effect sizes.

Receiver operating characteristic (ROC) curve analysis was performed to evaluate discriminatory performance. Area under the curve (AUC) values were calculated using the pROC package in R. Ninety-five percent confidence intervals were not calculated, as AUCs were derived from unsupervised RF models. Sensitivity and specificity were determined from ROC curves.

Clinical variables and outcomes were compared between groups using Wilcoxon rank-sum tests for continuous variables and Fisher’s exact tests for categorical variables. Statistical significance was defined as p < 0.05.

### Pathway Analysis

Associations between miRNAs and proteins with canonical pathways were assessed using miEAA (miRNA Enrichment and Annotation)^21,22^ and Ingenuity Pathway Analysis (IPA). Over-representation analysis was performed for miRNAs and proteins. Differential expression compared the secretome of pediatric DCM patients with poor outcomes vs those who recovered, using fold-change thresholds (FC > 0.5 or FC < –0.5). To increase the likelihood of identifying potential pathways for evaluation, we used a cutoff of p-value < 0.05 (–log_10_(p-value) > 1.3).

## RESULTS

### Cohort characteristics

Serum samples were collected from children diagnosed with dilated cardiomyopathy (DCM) (n = 70) at Children’s Hospital Colorado, Anschutz Medical Campus, Aurora, and the Pediatric Cardiomyopathy Registry (PCMR).^19^ Clinical characteristics of the cohort are presented in Table 1. DCM samples were obtained from patients under 21 years of age.

**Table 1:** Demographics characteristics and outcomes of subjects.

|  | Poor outcome | Recovered | Stable |
| --- | --- | --- | --- |
| *N* | 30 | 19 | 21 |
| Age, years, mean (IQR) | 9.9(13.0) | 3.3(1.7)* | 10.6(12.2) |
| Sex |  |  |  |
| Female, % | 55.2 | 46.7 | 66.6 |
| Male, % | 44.8 | 53.3 | 33.4 |
| Etiology (%) |  |  |  |
| iDCM | 96.6 | 89.3 | 100 |
| MC | 3.4 | 10.7 |  |
| Outcome (%) |  |  |  |
| Transplant | 69 | NA | NA |
| LVAD | 24.1 |  |  |
| Death | 6.9 |  |  |
| EF, %, mean±SD | 24.99±11.8 | 30.15±11.89 | 26.3±8.8 |
| Medications (%) |  |  |  |
| PDE3i, % | 42.9 | 38.5 | 5 |
| Digoxin, % | 20.7 | 30.8 | 40 |
| ACEi | 31.0 | 76.9 | 90.5 |
| Beta-blockers | 24.1 | 38.5 | 75 |
| Diuretics | 33.3 | 61.5 | 35 |

Abbreviations: iDCM, idiopathic dilated cardiomyopathy; MC, myocarditis; LVAD, left ventricular assist device, EF, ejection fraction; PDE3i, phosphodiesterase-3 inhibitor; ACEi, angiotensin convert enzyme inhibitor. \* $P = 0.0040$ for poor outcome and $P = 0.0090$ for stable.

Primary clinical outcomes were assessed one year after enrollment and classified as follows: poor outcome (n = 30), defined as requiring heart transplant (69%), mechanical circulatory support (24.1%), or death (6.9%); recovery (n = 19), defined as normalization of ventricular size and function; or stable (n = 21), defined as no disease progression and clinical compensation under medical therapy. Poor outcome and stable groups were generally age-matched; however, the recovery group was significantly younger.

There was approximately equal representation of male and female participants in the poor outcome and recovery groups, but not in stable patients where 66.6% was female while 33.4% was male. Most common disease etiology in both outcome groups was idiopathic DCM (96.6% in the poor outcome group and 89.3% in the recovery group). Myocarditis was also diagnosed in a subset of patients (3.4% in the poor outcome group and 10.7% in the recovery group).

LV ejection fraction (EF) values were derived from echocardiograms performed at the time of hospital admission. All patients presented with reduced EF, consistent with heart failure and subsequently received standard combination therapy for heart disease, including phosphodiesterase (PDE3) inhibitors, digoxin, angiotensin converting enzyme (ACE) inhibitors, beta-blockers, and/or diuretics.

### Identification of differentially expressed circulating miRNAs in DCM pediatric patients with poor outcomes vs recovery

To determine whether circulating miRNAs can predict recovery in children with DCM, total RNA was extracted from serum samples obtained at the time of presentation with acute heart failure. miRNAs were identified by RNA-seq. Poor outcomes or recovery were clinically determined one year after blood collection or at the time the outcome was met.

Differentially expressed miRNAs were identified using random forest (RF) analysis. The relative importance of candidate miRNAs, ranked by Mean Decrease in Accuracy and Gini, is shown in Figure 1A. Among these, hsa-miR-335-3p, hsa-miR-323a-3p, and hsa-miR-874-3p were the top three features contributing to classification. To evaluate the discriminatory capacity of these selected features, multidimensional scaling (MDS) based on the RF proximity matrix constructed from the top three miRNAs was performed. The MDS plot demonstrated clear separation between pediatric DCM patients with poor outcomes (n = 30) and those who recovered (n = 19) (Figure 1B), indicating distinct expression patterns captured by this three-miRNA panel.

**Figure 1.**
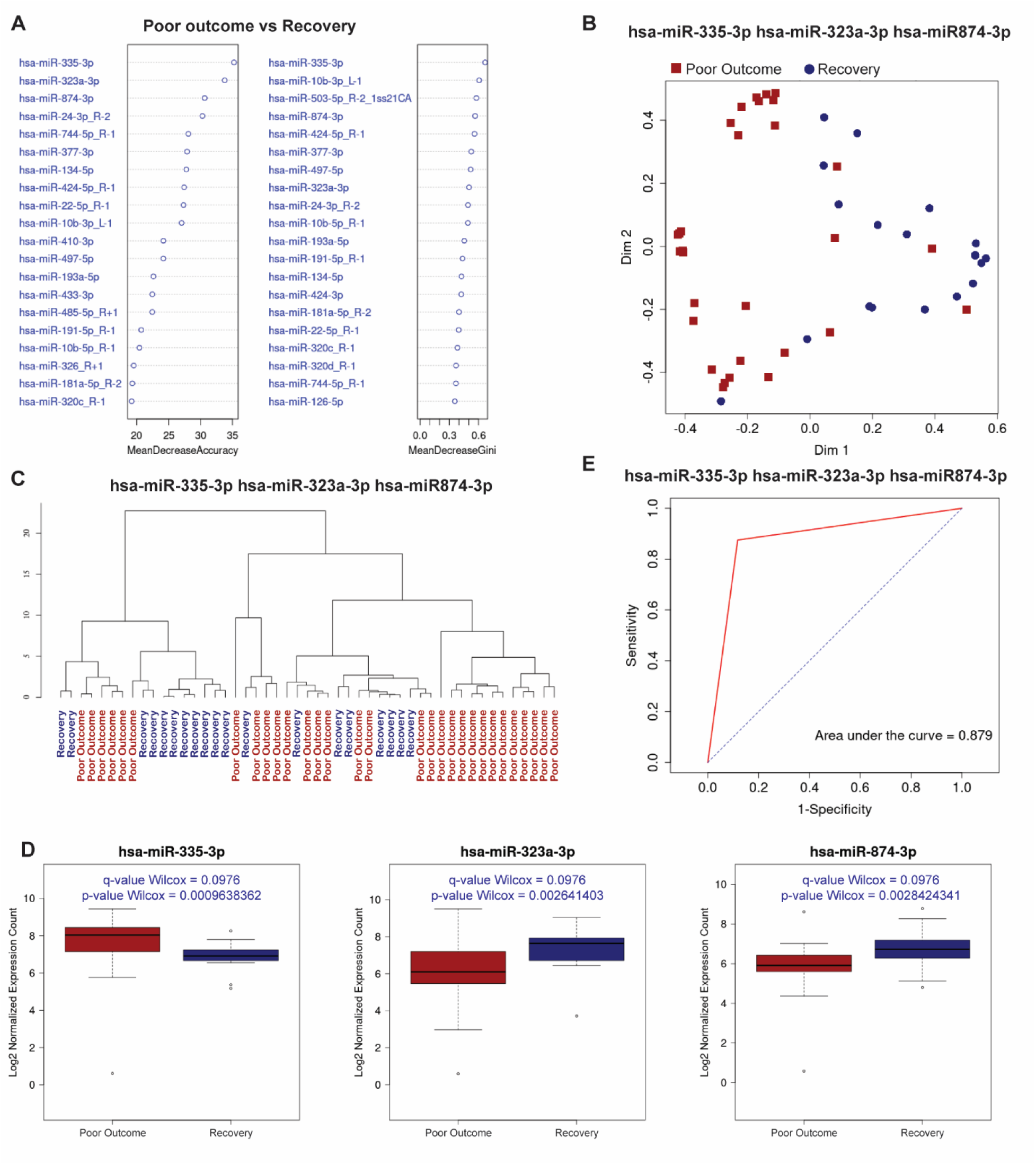
Circulating microRNAs (miRNAs) can differentiate heart failure outcomes in pediatric patients. (A) Random Forest (RF) variable importance plot highlights the contribution of each miRNA to group classification. (B) Random Forest (RF) analysis identified miR-335-3p, miR-323a-3p, and miR-874-3p as key circulating miRNAs that distinguish patients with poor outcomes (n = 30, red squares) from those who recovered (n = 19, blue circles). All samples are from the time of presentation with acute heart failure. (C) Hierarchical clustering based on the top three miRNAs shows clear separation between outcome groups. (D) Box plots display significant differences (q-value < 0.1) in serum levels of the three miRNAs between poor outcome and recovered patients. (E) ROC analysis using these top miRNAs yields an area under the curve (AUC) of 0.879, indicating strong discriminative potential for predicting poor outcomes.

Consistently, hierarchical clustering based on these top three miRNAs further supported separation between outcome groups (Figure 1C). Box plot analysis demonstrated statistically significant differences in expression levels between the two groups (Figure 1D). A receiver operating characteristic (ROC) curve generated from the three-miRNA panel yielded an area under the curve (AUC) of 0.879, indicating strong discriminative performance within this cohort for distinguishing pediatric DCM patients with poor outcomes from those who recovered (Figure 1E). The odds ratios for hsa-miR-335-3p, hsa-miR-323a-3p, and hsa-miR-874-3p were 1.623 (p = 0.039), 0.595 (p = 0.011), and 0.632 (p = 0.067), respectively. These results indicated that hsa-miR-335-3p is associated with increased odds of the event, whereas hsa-miR-323a-3p and hsa-miR-874-3p are associated with decreased odds. Based on feature selection metrics in our bioinformatic model, hsa-miR-335-3p and hsa-miR-323a-3p were classified among the strongest predictors, whereas hsa-miR-874-3p showed a moderate effect.

### Distinct subset of circulating proteins in DCM patients with poor outcomes vs recovery

Circulating protein profiles were identified from the serum samples of pediatric DCM patients. Like the miRNA findings, protein profiles from patients who eventually recovered were compared to those who had a poor outcome. The variable importance plot (Figure 2A), derived from RF analysis, ranked individual proteins according to their ability to differentiate between the two groups. RF analysis showed that CXCL12, ADGRF5, and COL2A1 could distinguish patients in the two groups (Figure 2B), which was further confirmed by hierarchical clustering (Figure 2C). Box plots illustrated the significant differences in expression of these proteins between the groups: CXCL12 was elevated in the poor outcomes group compared to patients who eventually recovered, while ADGRF5 and COL2A1 were decreased (Figure 2D). The ROC curve (Figure 2E) demonstrated an AUC of 0.865, consistent with high discriminative performance of these three proteins in distinguishing patients with poor outcomes from those who recovered. The odds ratios for CXCL12, ADGRF5, and COL2A1 were 15.460 (p = 4.03 × 10⁻⁶), 0.06216 (p = 1.7 × 10⁻⁴), and 0.419 (p = 5.6 × 10⁻⁴), respectively. According to our bioinformatic models, these values indicated strong predictors. Specifically, the high odds ratio for CXCL12 suggested an increased likelihood of poor outcomes, consistent with its upregulation in this group. Conversely, the lower odds ratios for ADGRF5 and COL2A1, which are downregulated in poor outcomes, suggested a protective effect and association with recovery.

**Figure 2.**
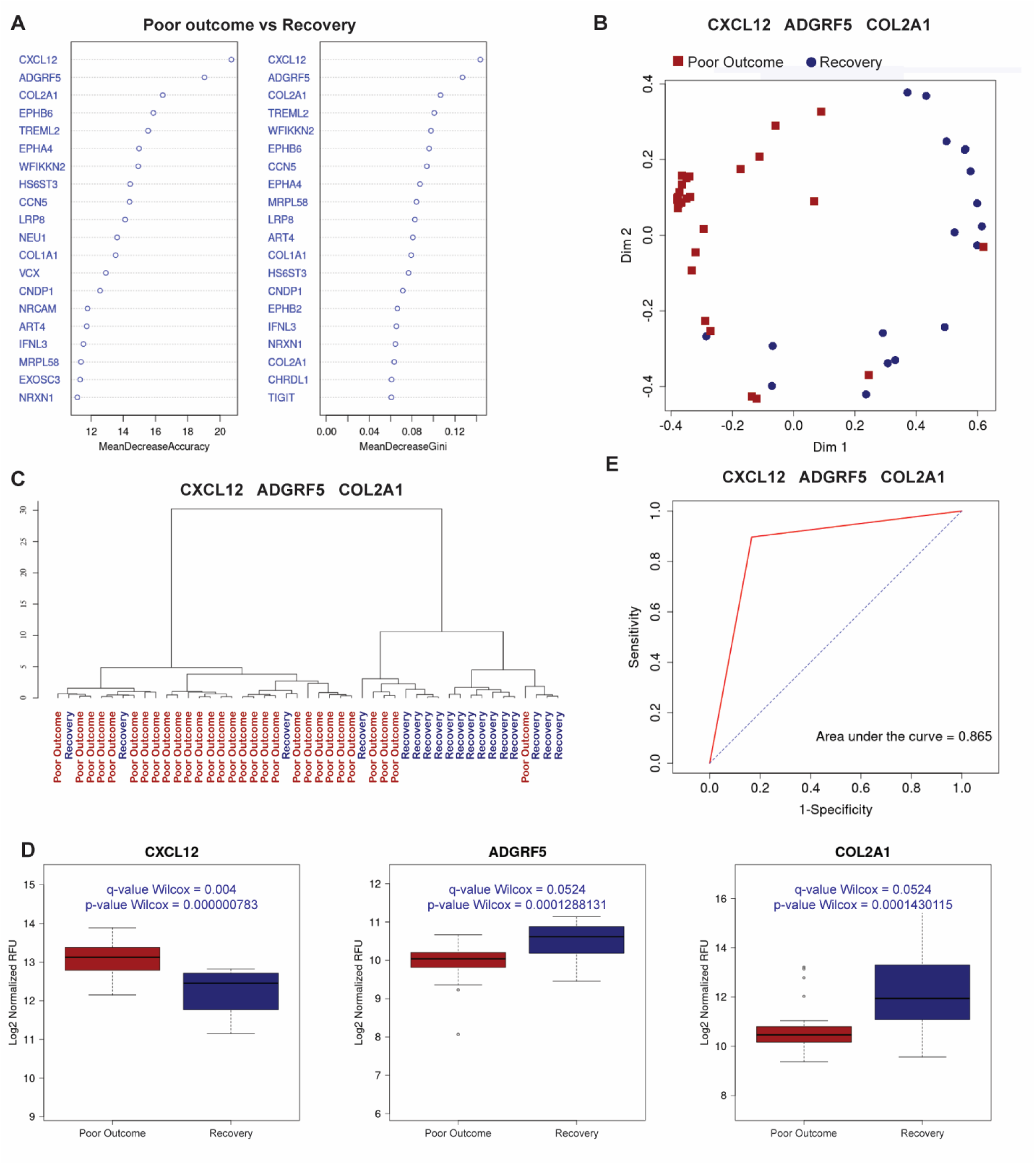
Circulating proteins earlier differentiate outcomes in pediatric acute heart failure. (A) RF variable importance plot showing the mean decrease in accuracy and mean decrease in Gini index for each circulating factor. The further a protein is to the right on the plot, the greater its contribution to distinguishing between the two clinical outcome groups. Features (aptamers) are ranked top-to-bottom by their overall importance in the model. (B) RF analysis identified circulating proteins, including CXCL12, ADGRF5, and EPHB6, distinguish patients with poor outcomes (red squares, n = 29) from those who recovered (blue circles, n = 18) at the time of presentation with acute heart failure. (C) Hierarchical clustering using the top three ranked proteins (CXCL12, ADGRF5, and EPHB6) shows clear separation between patients with poor outcomes and those who recovered, supporting their potential as predictive biomarkers. (D) Box plots show significant differences (adjusted p-value < 0.1) in serum protein levels of EPHB6, ADGRF5, and CXCL12 between pediatric heart failure patients with poor outcomes and those who recovered. (E) ROC analysis using the top three proteins yielded an area under the curve of 0.865, indicating good discriminative power for predicting poor outcomes.

### Combined analysis of circulating miRNAs and proteins is associated with improved discriminatory performance

To evaluate whether integrating circulating miRNAs and proteins improves disease outcome prediction, we performed a combined analysis using the top three miRNAs and the top three proteins identified by RF multidimensional scaling from samples at the time of presentation with acute HF. Only samples evaluated in both analyses were used in these investigations. RF analysis (Figure 3A) demonstrated that when hsa-miR-335-3p, hsa-miR-323a-5p, hsa-miR-874-3p, CXCL12, ADGRF5, and COL2A1 were analyzed together, groups were better defined. This finding was confirmed by HC analysis (Figure 3B). The RF variable importance plot (Figure 3C) highlighted the contribution of each miRNA and protein as a biomarker. Moreover, the combined analysis revealed a robust ROC curve with an AUC of 0.92 (Figure 3D), indicating that the integration of these six circulating factors provides a stronger predictive model for DCM outcome than individual factors alone. Using regression analysis, our results suggest a positive correlation between COL2A1 or ADGRF5 and hsa-miR-323a-3p and hsa-miR-874-3p. CXCL12, although not statistically significant, shows a negative correlation with hsa-miR-323a-3p and hsa-miR-874-3p (Figure 3E). This suggests that the distinctive signature identified in this acute heart failure scenario may be the result of similar biological processes and may aid in understanding mechanisms associated with pediatric DCM.

**Figure 3.**
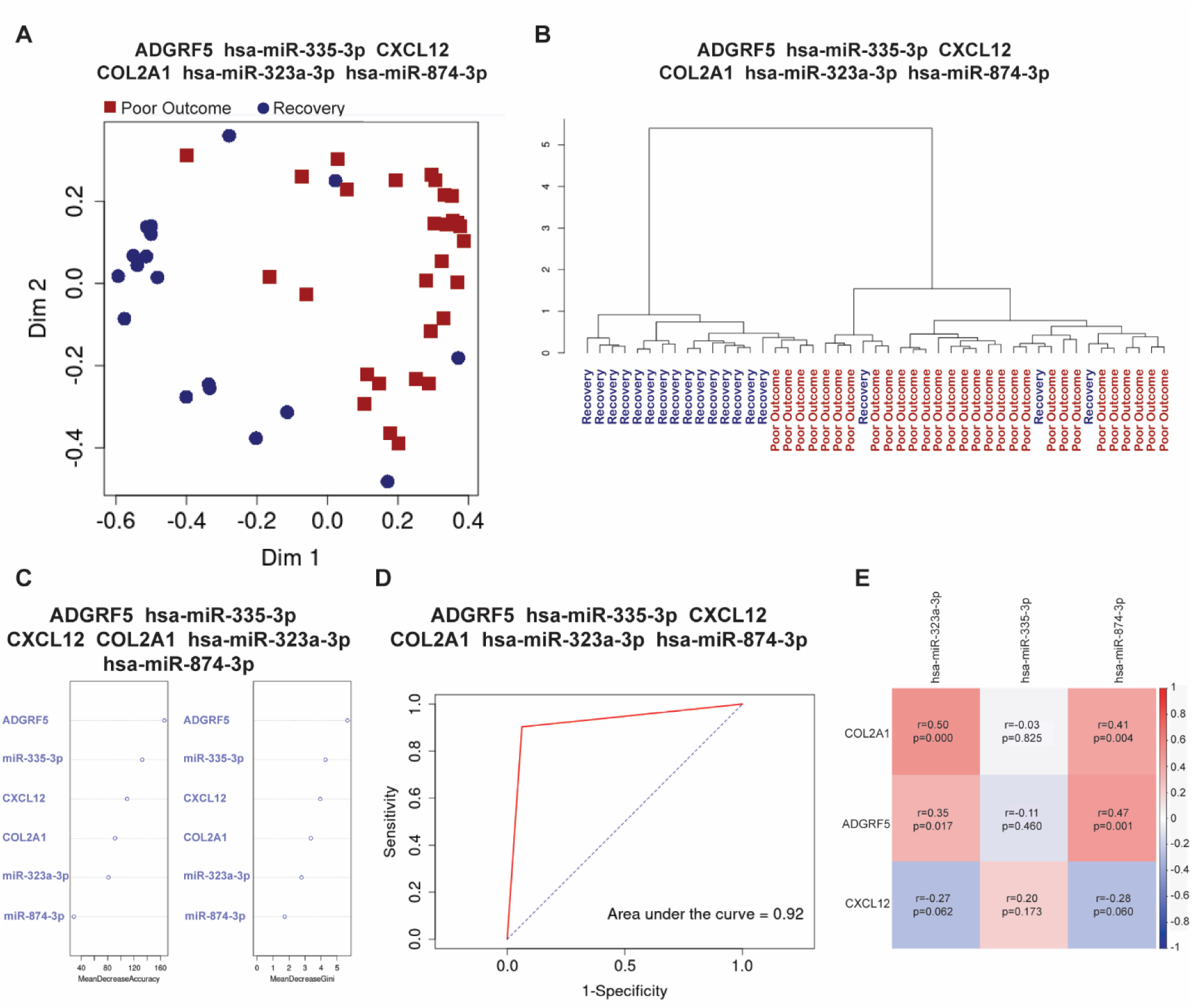
Combined analysis of top 3 circulating miRNAs and proteins. Combined analysis of the top three circulating miRNAs and proteins shows improved predictive performance when these biomarkers are used together. (A) Multidimensional scaling (MDS) based on RF analysis illustrates the estimated proximity matrix, (B) while the RF variable importance plot highlights the contribution of each miRNA and protein as a biomarker. (C) HC of the top three miRNAs and proteins demonstrates clear separation between outcome groups. (D) ROC analysis combining these biomarkers yields an AUC of 0.92, indicating high discriminative potential for predicting poor outcomes. (E) Regression analysis of the top 3 miRNAs and proteins shows significant relationships among the factors identified as discriminative.

### Paired analysis suggests that COL2A1, CXCL12, and ADGRF5 may be predictive biomarkers of outcomes over time

We next evaluated circulating miRNAs and proteins that can differentiate patients at the time of outcomes. A new subset of miRNAs (hsa-miR-342-3p, hsa-miR-222-3p-R+3, hsa-miR-26a-5p) and proteins (FABP4, ZP4, TTC9) was identified when comparing outcome samples from patients with poor outcomes vs those who recovered, suggesting that the secretome is malleable and changes as disease progresses.

We then tested if factors identified at the time of enrollment distinguished groups at the time of outcomes. The top three miRNAs (hsa-miR-335-3p, hsa-miR-323a-3p, hsa-miR-874-3p, Figure 4B.1) did not distinguish the two groups at outcomes, whereas the top three proteins (COL2A1, CXCL12 and, ADGRF5) (Figure 4B.2) showed a persistent ability to distinguish the groups, albeit with a lower accuracy, as reflected by the AUC of 0.675 (Figure 4D).

**Figure 4.**
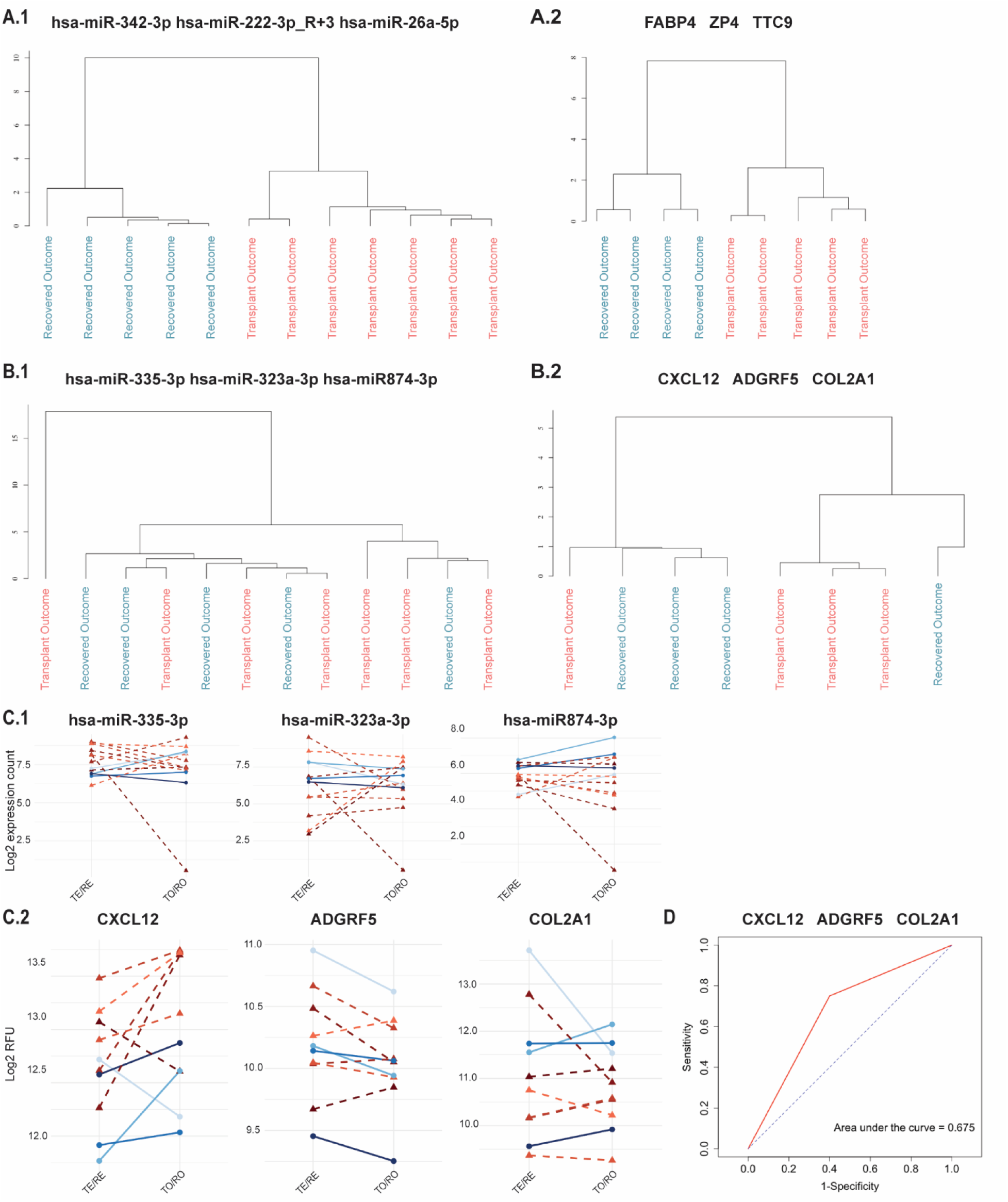
Longitudinal analysis of circulating factors reveals differences between the acute heart phase and the outcome. (A) HC analysis of circulating miRNAs (A.1) and proteins (A.2) demonstrates distinct subsets between patients who progressed to transplant and those who recovered at least 6 months after acute heart failure presentation. (B) HC analysis of the top 3 miRNAs (B.1) and proteins (B.2) at the time of acute heart failure shows that miR-335-3p, miR-323a-3p, and miR-874-3p do not separate the groups in follow-up samples. (C) Paired analysis of the same patients over time, at the time of heart failure (TE/RE) and after the outcome, at least 6 months later (TO/RO), demonstrates longitudinal variation of miRNAs (miR-335-3p, miR-323a-3p, miR-874-3p, C.1) and proteins (CXCL12, ADGRF5, COL2A1, C.2). (D) ROC curve analysis for CXCL12, ADGRF5, and COL2A1 in the follow-up samples indicates that this protein subset continues to have predictive value during the outcome phase. Reddish dashed lines represent the poor outcome group, and bluish solid lines represent the recovery group. TE/RE indicate transplant (or poor outcome) enrollment / recovered enrollment; TO/RO indicate transplant (or poor outcome) outcome / recovered outcome.

Analysis of miRNAs over time (Figure 4C.1) showed that their levels are variable as time progresses, suggesting their predictive ability is diminished longitudinally. In contrast, the predictive ability of COL2A1 and CXCL12 seems constant over time (Figure 4C.2). However, the limited longitudinal analysis, due to small size of sample, makes it difficult to establish robust associations over time.

Additionally, we evaluated levels of the top 15 miRNAs and proteins (as determined by statistical significance (q-value Wilcoxon < 0.1)) over time (samples were collected at enrollment and 6 to 12 months after). Expression levels of most miRs and proteins are not stable over time (data not shown), except for hsa-miR-10b-5p_R-1, hsa-miR-181-5p_R-2, MARCKSL1, ART4 (Figure S1). As the number of paired samples is significantly lower than the overall analysis, these results need to be further verified.

### Enrichment analysis provides insight into the mechanisms that differentiate poor outcome from recovered patients

We next evaluated the predicted biological significance of differentially expressed miRNAs at the time of presentation with acute heart failure (p-value Wilcoxon < 0.05 and fold change thresholds of > 0.5 for upregulated miRNAs and < −0.5 for downregulated miRNAs). Out of 459 miRNAs detected by miRNA sequencing, 75 had p-values below 0.05. Among these, 11 miRNAs showed a fold change > 0.5 and were considered upregulated in the poor outcome group, while 48 showed a fold change < −0.5 and were considered upregulated in the recovery group (Figure 5A).

**Figure 5.**
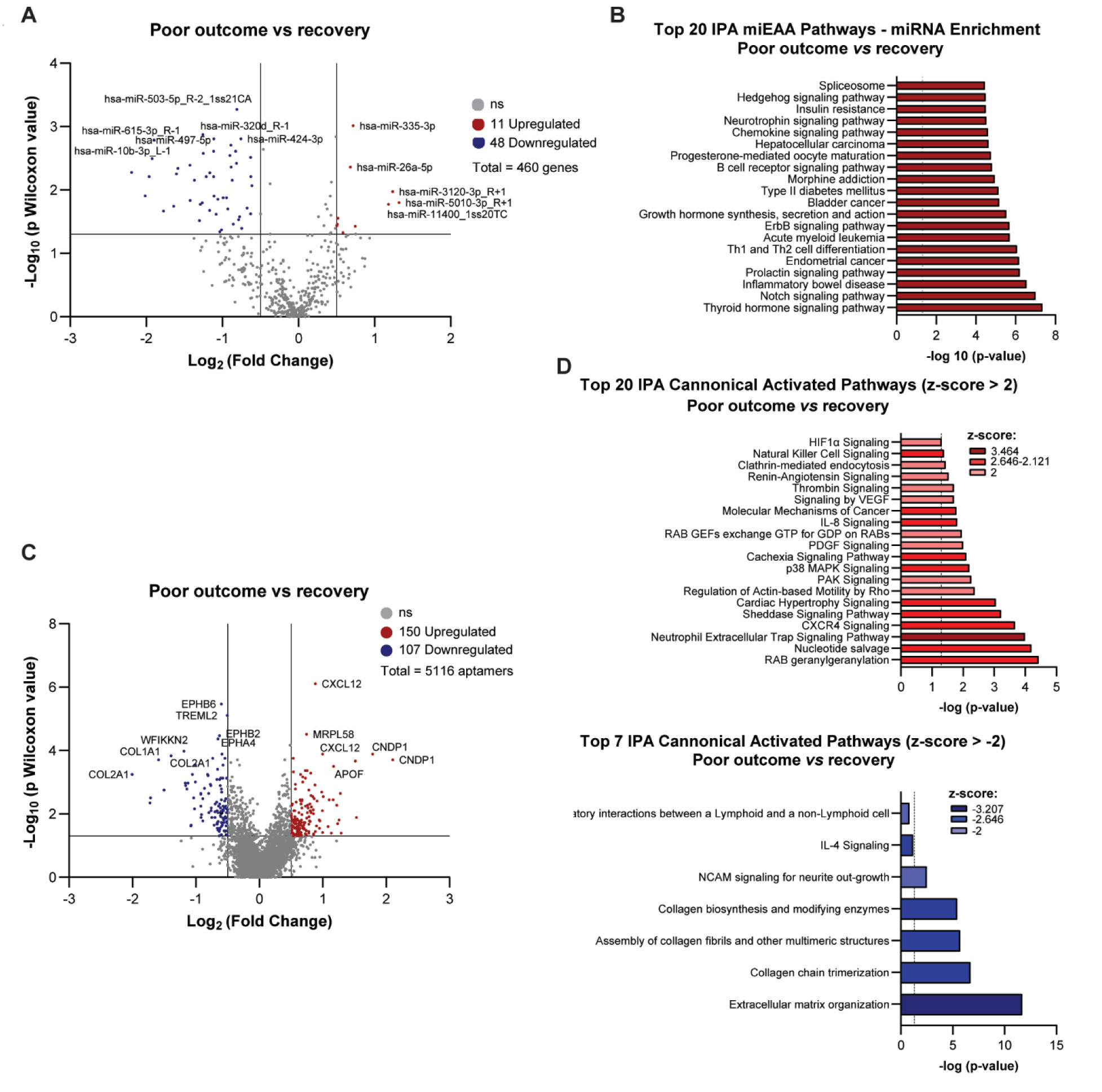
Secretome analysis reveals distinct miRNA and protein signatures and canonical pathway activation between pediatric DCM patients with poor outcomes and those who recovered. Differential expression analysis was performed by comparing the secretome of DCM pediatric patients with poor outcomes to those who recovered cardiac function (defined by fold change thresholds: > 0.5 or < –0.5 and –Log_10_ p-value > 1.3). (A) Volcano plot showing miRNAs significantly upregulated and downregulated in the poor outcome group compared to patients who eventually recovered ventricular function. (B) Top 20 predicted miEAA pathways predicted to be altered in response to dysregulated miRNAs. (C) Volcano plot showing proteins significantly upregulated and downregulated in the poor outcome group compared to the recovery group. (D) Canonical pathway analysis using Ingenuity Pathway Analysis (IPA). The top 20 enriched pathways with z-score > +2 represent pathways activated in the poor outcome group, while the top 7 pathways with z-score < –2 represent pathways downregulated in the poor outcome group. Enrichment significance is shown as –log (p-value), with a threshold of >1.3 (Fisher’s exact test). The intensity of red or blue shading indicates the predicted activation or inhibition state of the pathway, respectively.

Using miEAA pathways, the enrichment analysis of differentially expressed miRNAs (Figure 5B) revealed several key signaling pathways, which can be broadly grouped into immune/inflammatory (Th1 and Th2 cell differentiation, chemokine signaling, B cell receptor signaling, and inflammatory bowel disease), hormonal/metabolic (thyroid hormone signaling, prolactin signaling, growth hormone pathways, insulin resistance, and type II diabetes mellitus), developmental and growth signaling (ErbB, Notch, and Hedgehog), cancer-related (multiple cancer types, neurotrophic/stress responses neurotrophin signaling, morphine addiction), and RNA regulatory processes (Splicesome).

Applying the same threshold as for miRNAs, the biological significance of proteins was investigated. Out of 5,116 aptamers detected by SomaScan^®^, 626 had p-values < 0.05. Among these, 150 aptamers showed a fold change > 0.5 and were considered upregulated in the poor outcome group, while 107 aptamers showed a fold change < −0.5 and were considered upregulated in the recovery group (Figure 5C).

Pathway analysis, using the IPA platform, revealed that pediatric DCM patients with poor outcomes (Figure 5D, red bar graph) displayed a robust inflammatory component, characterized by activation (z-score > 2) of pathways including NET signaling, CCR4 signaling, p38 MAPK signaling, IL-8 signaling, natural killer cell signaling, thrombin signaling, and cachexia signaling. Additionally, this group showed enrichment of cardiovascular and stress response pathways (cardiac hypertrophy signaling and renin–angiotensin signaling), cytoskeletal dynamics (regulation of actin-based motility by Rho and PAK signaling), vesicle trafficking/endocytosis (RAB signaling and clathrin-mediated endocytosis), and growth factor/oncogenic pathways (PDGF Signaling, CXCR4 Signaling, Sheddase Signaling Pathway, Molecular Mechanisms of Cancer).

In contrast, enrichment analysis in the recovery group (from pathways predicted to be downregulated in the poor outcome group, with a z-score < −2) revealed a distinct signature dominated by ECM and collagen remodeling pathways (Figure 5D, blue bar graph). Importantly, although not statistically significant, IL-4 signaling emerged as a potential driver of a Th2-type inflammatory response, which may contribute to the observed ECM/collagen remodeling in these patients.

Together, the integration of miRNA and proteomic enrichment analyses demonstrates strong convergence on immune/inflammatory and cardiovascular remodeling mechanisms, while also revealing additional regulatory processes.

### A distinct miRNA and protein signatures were observed between patients who had poor outcomes and those with stable DCM

The circulating miRNA and protein profiles were also investigated in stable patients. The importance (IMP) analysis ranked the top three most relevant miRNAs (Figure S2.A). Random Forest (Figure S2.B) identified miR-548av-5p, miR-215-5p_R-1, and miR-7703-p5_1ss3GT as key miRNAs differentiating stable from poor outcomes, which was further supported by hierarchical clustering (Figure S2.C). These miRNAs are statistically different (q-value Wilcoxon < 0.1) (Figure S2.D) between the groups, yielding a reasonably good ROC curve fit (AUC = 0.819) (Figure S2.E).

Similarly, analysis of circulating proteins (Figure S3), using the same approach, identified three potential candidates that statistically distinguish (q-value Wilcoxon < 0.1) severe DCM from stable DCM with a ROC curve showing an AUC of 0.899: NECAP2, UBQLN2, and GGA1. All three proteins were found to be increased in patients with poor outcomes compared to those with stable disease.

### Circulating miRNAs and proteins show an ability to differentiate patients with stable DCM from those who recovered

Using a similar approach, we compared stable DCM patients with patients who eventually recovered. miRNA analysis identified miR-548av-5p_R+4, miR-22-3p, and miR-548I_R-1 as the top differentiating miRNAs when comparing stable and recovered patients. The ROC curve showed an AUC of 0.799, suggesting miRNAs can separate both groups (Figure S4).

Additionally, protein analysis (Figure S5), using RF, identified three top candidates, RILPL1, CSF1R, and CARNMT1, which show a strong discriminative performance to separate stable patients from patients who eventually recovered cardiac function (AUC = 1). Notably, all three proteins were significantly increased in the recovery group compared with stable DCM.

## DISCUSSION

The exploration of diverse biomolecules as molecular biomarkers through multi-omics analysis has expanded, offering opportunities to improve disease prevention, diagnosis, prognosis, and monitoring, while also providing insights into pathogenic mechanisms and facilitating the development of targeted therapies. These approaches hold great promise for bridging the research gap in pediatric DCM.

This study provides a comprehensive secretome analysis to identify circulating factors that may predict clinical outcomes in pediatric patients with acute heart failure due to DCM. We used miRNA-sequencing and SomaScan^®^ proteomics to define miRNA and protein signatures characteristic of patients with severe DCM who progressed to poor outcomes, remained stable, or recovered. This approach offers a unique perspective on potential biomarkers in this patient population. We found that circulating miRNAs and proteins robustly discriminated patient outcomes in children with DCM. Importantly, these differences were detectable at the time of the first blood collection during the initial hospital visit, shortly after symptom onset. This finding suggests that circulating mediators released during acute heart failure may serve as early predictors of long-term outcomes.

We identified a subset of miRNAs (hsa-miR-335-3p, hsa-miR-323a-3p, hsa-miR-874-3p) and proteins (CXCL12, ADGRF5, COL2A1) that distinguished children with DCM who had poor outcomes (i.e., required transplant, mechanical assisted device, or died) from those who eventually recovered. Additionally, levels of miR-548av-5p_R+4, miR-215-5p_R-1, and miR-7703-5p_1ss3GT, and the proteins NECAP2, UBQLN2, and GGA1, were altered between patients with poor outcomes and those who remained stable. Moreover, miR-548av-5p_R+4, miR-22-3p, miR-548I_R-1, and the proteins RILPL1, CSF1R, and CARNMT1 distinguished stable patients from those who recovered. However, protein analysis included only six stable patients, making it difficult to determine with confidence whether these proteins can reliably serve as biomarkers to distinguish stable from severe or recovered DCM.

When we performed combined analyses, the top three miRNAs and top three proteins distinguishing poor outcomes from recovery demonstrated a more robust separation than when assessed individually. This suggests that these six molecules could be used in combination to more accurately predict clinical outcomes. To our knowledge, these data provide the first evidence that integrating miRNA and protein signatures can help determine prognosis in pediatric DCM.

Pathway enrichment analysis revealed distinct biological profiles across groups. The poor-outcome group was enriched in pathways related to inflammatory responses, cardiac remodeling, cytoskeletal dynamics, and vesicle trafficking/endocytosis. In contrast, the recovery group showed predominant enrichment in ECM and collagen remodeling pathways, underscoring divergent mechanisms driving recovery vs disease progression.

In this study, CXCL12 was identified as a circulating factor elevated in the poor-outcome group, suggesting that an inflammatory component may contribute to disease progression. While it can recruit beneficial stem cells for cardiac repair, CXCL12 also directly contributes to cardiac injury by promoting apoptosis, hypertrophy, and inflammation in cardiomyocytes, ultimately leading to worse outcomes in heart failure. Its specific role in DCM remains complex and not fully understood, with studies suggesting it may serve as both a biomarker and a potential therapeutic target, particularly in processes such as fibrosis.^23^

In coronary artery disease (CAD), CXCL12 has been linked to adverse outcomes. Chang et al. first reported that higher serum CXCL12 levels predicted major adverse clinical events within 30 days of acute myocardial infarction (AMI).^24^ Subsequent studies further supported a positive correlation between elevated circulating CXCL12 and increased short- and long-term cardiovascular risk.^25–27^ However, contrasting evidence showed that higher CXCL12 levels could also predict lower future cardiovascular events.^28^ Thus, the relationship between circulating CXCL12 and adverse cardiovascular outcomes in CAD remains controversial.

Interestingly, we also found a negative correlation between CXCL12 and miR-874-3p. miR-874-3p was enriched in the recovery group, in contrast to the elevated CXCL12 levels observed in poor-outcome patients. Previous work has shown that miR-874-3p regulates inflammation in alveolar epithelial cells by targeting EGR3/NFκB,^29^ attenuates macrophage-mediated inflammation in intracerebral hemorrhage^30^ and, notably, suppresses CXCL12 expression to promote angiogenesis and reduce inflammatory responses in ischemic stroke.^31^ These results suggest that higher miR-874-3p levels may aid in improving disease and may suggest that miR-874-3p can contribute to lower CXCL12 levels.

Additionally, we observed increased levels of miR-335-3p in the poor-outcome group compared to the recovery group. This miRNA has been reported to regulate cardiac mesoderm and progenitor cell differentiation through activation of WNT and TGF-β signaling pathways.^32^ We have recently reported that WNT signaling is altered in the pediatric DCM heart and can be a key driver of cardiac remodeling, stiffness, and disease progression.^33,34^ This suggests that the increase in miR-335-3p may contribute to activation of detrimental signaling pathways.

On the other hand, elevated circulating levels of COL2A1 were found in the recovery group, distinguishing them from the poor-outcome group. This biomarker suggests strong involvement of extracellular matrix and collagen remodeling processes. Interestingly, there is minimal fibrosis in explanted hearts of children with DCM.^35,36^ A strong signature of collagen remodeling processes may suggest that an impairment of fibrotic responses may contribute to poor outcomes. Although highly speculative, the faster disease progression in children may be associated with the inability to promote compensatory remodeling, such as lack of cardiomyocyte hypertrophy and minimal fibrosis.^33,35,36^

Importantly, COL2A1, a major component of cartilage, intervertebral discs, and the vitreous humor, can be released into circulation during turnover, injury, or inflammation.^37^ COL2A1 has shown a strong association with WNT/β-catenin signaling. In human articular cartilage explants, activation of the canonical WNT/β-catenin pathway (e.g., using LiCl) after mechanical injury increased COL2A1 and aggrecan mRNA expression.^38^ In contrast, in chick sternal chondrocytes, overexpression of Wnt8c, Wnt9a, or β-catenin inhibited COL2A1 transcription while promoting hypertrophic markers such as Col10a1 and Runx2.^39^ Furthermore, loss of β-catenin in aortic valve interstitial cells promoted chondrogenic differentiation, leading to increased SOX9 nuclear localization and enhanced expression of *Col2a1, Acan, Col10a1*. This phenotype, characterized by excessive chondrogenic proteoglycan accumulation and disrupted ECM stratification, is typical of myxomatous valve disease.^40^ Additionally, since COL2A1 is involved in cartilage and extracellular matrix biology, and miR-323a-3p (increased in recovery group) has been implicated in fibrosis^41,42^ and epithelial–mesenchymal transition,^43^ there may be indirect regulatory relationships. Specifically, miR-323a-3p could modulate upstream transcription factors and signaling molecules (e.g., SMAD2, TGFA), which in turn may influence COL2A1 expression.

Interestingly, the protein ADGRF5, a protein from the G-protein coupled receptor (GPCR) family, was found to be significantly increased in the recovery group compared to the poor-outcome group. This protein was recently characterized as a cardiomyocyte (CM)-specific factor involved in maintaining cardiac homeostasis. One preliminary study showed that ADGRF5 CM-specific knockout mice (F5cmKO) developed cardiac dysfunction, maladaptive remodeling, and increased mortality over time, even in the absence of pathological insult, supporting a homeostatic role for ADGRF5 in cardiomyocytes.^44^ These findings suggest that augmenting ADGRF5 may be a novel mechanism to improve heart function.

To confirm the robustness of the biomarker panel, this study also analyzed follow-up samples. These analyses showed that 6–12 months after the initial blood collection, other circulating factors better separate samples based on outcomes. hsa-miR-342-3p, hsa-miR-222-3p_R+3, hsa-miR-26a-5p, FABP4, ZP4, and TTC9 emerged as potential candidates for monitoring disease progression. This suggests that the secretome is malleable and secreted factors change as disease progresses. Although this may not be unexpected, it may limit the use of biomarkers at times other than presentation. We explored if the top six biomarkers from the time of enrollment in the study could inform prognosis over time. Our results are limited by the small sample size but indicate that miRNAs may not be ideal biomarkers later in the disease process. However, proteins may retain the ability to serve as good prognostic markers.

Altogether, our data provide a robust bioinformatic analysis identifying circulating factors - hsa-miR-335-3p, hsa-miR-323a-3p, hsa-miR-874-3p, CXCL12, ADGRF5, and COL2A1 - through an unbiased approach. These factors form a potential diagnostic “fingerprint” capable of informing prognosis at presentation, while also offering insights into the biological mechanisms underlying clinical trajectories, consistent with supporting literature. This work opens new fields of study for advancing our understanding of pediatric DCM.

## CONCLUSIONS

This work identifies a novel biomarker signature capable of predicting outcomes in children at the time of presentation with acute heart failure due to DCM. Additionally, it offers insights into pathways associated with poor prognosis and those important for recovery, which may guide the future identification of new therapeutic targets.

## LIMITATIONS

This study has some limitations. (1) There is a significant difference in the age of patients from the recovered and poor outcome groups, which may influence the levels of the factors identified as biomarkers. However, although younger age is associated with recovery, age alone has limited discriminatory power in prior pediatric DCM cohorts, suggesting that the circulating biomarker signatures identified here capture biological information beyond age-related risk. (2) The SomaScan® analysis in stable DCM patients was performed in only six subjects, limiting confidence in whether the subset of proteins identified as biomarkers for poor outcome vs. stable or stable vs. recovered can reliably distinguish stable from severe or recovered DCM. (3) The limited number of follow-up samples available to evaluate levels of circulating factors over time limits a clear understanding of the variation in the levels of these biomarkers. (4) Furthermore, the miRNAs identified in this study are different than what we previously identified as biomarkers of recovery. The current study is RNA-seq–based, whereas the prior study is array-based. In addition to a greater number of miRNAs identified by RNA-seq, arrays require a normalizer, which may artificially alter relative miRNA levels. Future studies including independent validation cohorts are needed to confirm the robustness of the biomarkers identified in this study. (5) Although a validation cohort is not available, the consistency of findings across multiple analytical approaches, such as Random Forest, Wilcoxon rank-sum testing, logistic regression, ROC analysis, and unsupervised hierarchical clustering, provides strong computational validation. These methods rely on different statistical assumptions, including nonlinear versus linear, parametric versus non-parametric, and supervised versus unsupervised. The agreement in results lowers the chances that our findings are influenced by model-specific bias or overfitting. Therefore, while experimental validation would be beneficial in future studies, the agreement from various methods supports the strength of the current results.

## ACKNOWLEDGMENTS

The authors thank Danielle Dauphin from the University at Buffalo, Jacobs School of Medicine and Biomedical Sciences, Buffalo, NY, for administrative support. We thank the Children’s Cardiomyopathy Foundation and the Kyle John Rymiszewski Foundation for their ongoing support of the Pediatric Cardiomyopathy Registry’s research efforts.

## DATA AVAILABILITY

The datasets supporting the current study including raw, processed and metadata of miRNA sequencing and SomaScan will be made publicly available in NCBI GEOdatabase upon acceptance of the manuscript for publication.

## NON-STANDARD ABBREVIATIONS AND ACRONYMS

ACEi: Angiotensin-converting enzyme inhibitor
ADGRF5: Adhesion G protein–coupled receptor F5
AMI: acute myocardial infarction
ART4: ADP-ribosyltransferase 4
AUC: Area under the curve
BNP: B-type natriuretic peptide
CAD: Coronary artery disease
CARNMT1: Carnosine N-methyltransferase 1
CCR4: C-C chemokine receptor type 4
CM: Cardiomyocyte
COL2A1: Collagen type II alpha 1 chain
CSFR1: Colony-stimulating factor 1 receptor
CXCL12: C-X-C motif chemokine ligand 12
CXCR4: C-X-C chemokine receptor type 4
ECM: Extracellular matrix
ERB: Erb-B2 receptor tyrosine kinase (HER2)
EGR3: Early growth response 3
ESC: European Society of Cardiology
FABP4: fatty acid-binding protein 4
GGA1: Golgi-associated, gamma adaptin ear containing, ARF binding protein 1
HC: Hierarchical clustering
iDCM: Idiopathic dilated cardiomyopathy
IPA: Ingenuity pathway analysis
IL-4: Interleukin 4
LV: Left ventricle
LV EF: Left ventricular ejection fraction
LV FS: Left ventricular fractional shortening
LVEDD: Left ventricular end-diastolic diameter
MAPK: Mitogen-activated protein kinase
MARCKSL1: MARCKS-like 1
MDS: Multidimensional scaling
miEAA: miRNA Enrichment and Annotation
miR: microRNA
miRNA: microRNA
NECAP2: NECAP endocytosis associated 2
NET: Neutrophil extracellular trap
NF-κB: Nuclear factor–kappa B
Notch: Notch signaling pathway
NT-proBNP: N-terminal pro–B-type natriuretic peptide
PAK: p21-activated kinase
PDE3i: Phosphodiesterase 3 inhibitor
PDGF: Platelet-derived growth factor
RAB: Ras-related protein Rab
RF: Random forest
Rho: Ras homolog family of small GTPases
RILPL1: Rab interacting lysosomal protein like 1
ROC: Receiver operating characteristic
SQS: SOMAscan quality statements
TE: transplant enrollment
TO: transplant outcome
RE: Recovery enrollment
RO: Recovery outcome
TTC9: Tetratricopeptide repeat domain 9
UBQLN2: Ubiquilin 2
WNT: Wnt signaling pathway
ZP4: Zona pellucida glycoprotein 4

